# XRCC1 mediates PARP1- and PAR-dependent recruitment of PARP2 to DNA damage sites

**DOI:** 10.1101/2024.05.14.594230

**Authors:** Xiaohui Lin, Kay Sze Karina Leung, Kaitlynn F. Wolfe, Brian J. Lee, Shan Zha

**Affiliations:** Institute for Cancer Genetics, Vagelos College for Physicians and Surgeons, Columbia University, New York City, NY 10032, USA; Columbia College, Columbia University, New York, NY 10027, USA; Department of Pathology and Cell Biology, Herbert Irvine Comprehensive Cancer Center, Vagelos College for Physicians and Surgeons, Columbia University, New York City, NY 10032, USA; Division of Hematology, Oncology and Stem Cell Transplantation, Department of Pediatrics, Vagelos College for Physicians and Surgeons, Columbia University, New York City, NY 10032, USA; Department of Immunology and Microbiology, Vagelos College for Physicians and Surgeons, Columbia University, New York City, NY 10032, USA

## Abstract

Poly-ADP-ribose polymerases 1 and 2 (PARP1 and PARP2) are crucial sensors of DNA-strand breaks and emerging cancer therapy targets. Once activated by DNA breaks, PARP1 and PARP2 generate poly-ADP-ribose (PAR) chains on themselves and other substrates to promote DNA single-strand break repair (SSBR). PARP1 can be activated by diverse DNA lesions, whereas PARP2 specifically recognizes 5’ phosphorylated nicks. They can be activated independently and provide mutual backup in the absence of the other. However, whether PARP1 and PARP2 have synergistic functions in DNA damage response remains elusive. Here, we show that PARP1 and the PAR chains generated by PARP1 recruit PARP2 to the vicinity of DNA damage sites through the scaffold protein XRCC1. Using quantitative live-cell imaging, we found that loss of XRCC1 markedly reduces irradiation-induced PARP2 foci in PARP1-proficient cells. The central BRCT domain (BRCT1) of XRCC1 binds to the PAR chain, while the C-terminal BRCT domain (BRCT2) of XRCC1 interacts with the catalytic domain of PARP2, facilitating its localization near the breaks. Together, these findings unveil a new function of XRCC1 in augmenting PARP2 recruitment in response to PARP1 activation and explain why PARP1, but not PARP2, is aggregated and hyperactivated in XRCC1-deficient cells.

## INTRODUCTION

Poly-ADP-ribose polymerase 1 (PARP1) and PARP2 are crucial sensors of DNA strand breaks and important targets of cancer therapy (1,2). Upon DNA strand breaks, PARP1 and PARP2 bind to DNA ends via their N-terminal domains, which initiates a series of allosteric changes that activate the C-terminal ADP-ribose transferase (ART) domains (3,4). PARP1 has five DNA binding moieties: three zinc finger domains, one Tyr-Gly-Arg (WGR) domain, and one BRCT domain, which collectively support nanomolar affinity binding and activation by various DNA strand breaks, from double-strand breaks (DSB) to single-strand nicks and gaps (4,5). In contrast, PARP2 possesses just two DNA binding domains: the unstructured N-terminal region (NTR) and the WGR domain, with only the latter required for break-induced activation of PARP2 (6). Consequentially, PARP2 is selectively activated by 5’phosphorylated nicks (4,7). The C-terminal catalytic (CAT) domain is highly conserved between PARP1 and PARP2. Activated PARP1 and PARP2 use nicotinamide adenine dinucleotide (NAD+) as a donor to add ADP-ribose (ADPr) to Serine, Aspartic Acid, and Glutamine residues of protein substrates, most notably themselves (1,2). PARP1 and PARP2 can extend the mono-ADPribosylation (MARylation) into complex straight or branched Poly-ADP-ribose (PAR) chains, termed PARylation. These negatively charged PAR chains facilitate chromatin relaxation and recruit other nucleic acidic binding proteins, including DNA repair factors and transcriptional regulators (8). DNA damage-induced PAR chains in physiological conditions are transient, reflecting both efficient repair and degradation by the abundant Poly-ADP-ribose Glycosylase (PARG). After single dose micro-irradiation with a 405nm laser, PAR foci form immediately and are mostly dissolved within 10 minutes (9,10). PARP1 and PARP2 can function independently, providing mutual backup if one is absent. Indeed, mice deficient for Parp1 or Parp2 were born alive, but the loss of both Parp1 and Parp2 caused embryonic lethality (11,12). However, in cells, PARP1 is much more abundant and robust than PARP2, accounting for > 80% of DNA damage induced PARylation (13). Given the lower prevalence of PARP2 and its overlapping DNA lesion specificity with PARP1, it is unclear whether and how PARP2 accesses DNA breaks in the presence of PARP1.

Among the proteins recruited by PAR chains to the DNA damage response is the X-ray repair cross-complementing group 1 (XRCC1), a molecular scaffold protein critical for single-strand DNA break repair (SSBR). XRCC1 is comprised of an N-terminal domain, a central BRCT domain (BRCT1), and a C-terminal BRCT domain (BRCT2). The BRCT1 domain of XRCC1 contains a phosphorylate binding pocket that can also bind to PAR(14), facilitating the recruitment of XRCC1 and its associated proteins to the site of damage (32–35). Notable cargos of XRCC1 include the DNA Polymerase β (Polβ), which binds to the N-terminal domain of XRCC1, and the DNA Ligase 3 (LIG3), which binds to the BRCT2 domain (15,16). In this regard, the XRCC1 is also essential for the protein stability of nuclear LIG3 (17). The linker between BRCT1 and BRCT2 contains multiple phosphorylation sites that are recognized by Polynucleotide kinase phosphatase (PNKP), Aprataxin (APTX), and Aprataxin and PNKP-Like Factor (APLF), which play critical roles in DNA end-processing and chromatin remodeling, respectively (18–20). Supporting the importance of XRCC1 and its partners in PAR-mediated SSBR, XRCC1 deficiency causes embryonic lethality in mice accompanied by toxic trapping and hyperactivation of PARP1 (21). Correspondingly, PARP1 deletion significantly alleviates the transcription and DNA repair defects observed in XRCC1-deficient cells (22,23), highlighting the feedback regulation between XRCC1 and PARP1. In this context, FDA-approved dual PARP1 and PARP2 inhibitors (PARPi) significantly delay PAR-dependent XRCC1 recruitment and cause continuous presence of PARP1, termed trapping, at the DNA damage sites (9,24). Loss of PARP1 decreases PARPi sensitivity by ∼10 folds, indicating the critical contribution of PARP1-trapping to the therapeutic effect of PARPi (9). In theory, unrepaired breaks accumulated in XRCC1-deficient or PARPi-treated cells should also recruit PARP2. Although FDA-approved PARPi causes allosteric trapping of PARP2 (10,25), hyperactivation of PARP2 has not been described for XRCC1-deficient cells.

While investigating the impact of FDA-approved PARPi on PARP2 recruitment and exchange, we identified two modes of PARP2 recruitment using quantitative live-cell imaging analyses: PARP1-independent and PARP1-dependent (10). The PARP1-independent pathway relies on the direct interaction between the PARP2 WGR domain with DNA breaks (6,10). The R140A mutation within the PARP2 WGR domain, impeding DNA binding (26), completely abolishes the break-induced PARP2 foci in PARP1-deficient cells (10). However, in the presence of PARP1, even PARP2-R140A, lacking DNA binding capacity, can still form robust foci. Moreover, PARPi, niraparib, largely abolished the PARP2-R140A foci in PARP2 proficient cells, indicating that it is the catalytic activity of PARP1 and the PAR chain, but not PARP1 protein itself, that facilitates the recruitment of PARP2-R140A (10). How could PAR promote PARP2 recruitment? One previous study suggested that the unstructured NTR of PARP2 can bind to PAR and is sufficient to recruit NTR-tagged GFP to the DNA damage sites (27). Yet, we and others found that in the context of full-length PARP2, loss of NTR did not significantly impact PARP2 recruitment to DNA damage sites (6,10), suggesting an alternative model exists. Here, we identify XRCC1 as a critical mediator for PARP1 and PAR-dependent recruitment of PARP2 to the vicinity of DNA breaks. In contrast to PARP1, which is hyperactivated and aggregates at the break site in *XRCC1 KO* cells, PARP2 foci are diminished in XRCC*1 KO* cells. Furthermore, we find that XRCC1 promotes the local recruitment of PARP2 by binding to PAR via its BRCT1 domain and PARP2 via its BRCT2 domain. Together, these results identified a previously unappreciated mechanism by which PARP1 promotes PARP2 local enrichment around breaks. These results explain the delayed PARP2 foci formation documented in previous studies and why only PARP1, not PARP2, is hyperactivated in XRCC1-deficient cells. They also predict that loss of PARP1 would compromise early PARP2 recruitments.

## MATERIALS AND METHODS

### Cell lines and cell culture

The TERT-immortalized human retinal pigment epithelial-1 (RPE-1) cell lines, including wild type (WT), *PARP1* knockout (KO), *PARP2* KO, *PARP1/2* double knockout (DKO), *XRCC1 KO* and *PARP1/XRCC1 DKO* variants, were generously provided by Dr. Keith W. Caldecott at the University of Sussex (28,29). The immortalized murine embryonic fibroblast (iMEFs) were generated in-house. Briefly, we transduced primary MEFs isolated from E13.5 embryos with retrovirus encoding SV40 large and small antigens as previously described (30). Primary MEFs were derived from timed breeding using a standard protocol. The *Xrcc1 KO* iMEFs were generously provided by Dr. Li Lan, who is currently at Duke University. Cells were cultured in DMEM medium (GIBCO, Cat. 12430062) supplemented with fetal bovine serum (FBS, 10% for RPE-1 cells and 15% for iMEFs), MEM non-essential amino acids (GIBCO, Cat. 11140050), 2 mM glutamine, 1 mM sodium pyruvate, and 50 U/mL penicillin/streptomycin (GIBCO, 15140122).

### Chemicals and antibodies

The PARP inhibitor niraparib (Selleckchem, S2741) was dissolved in dimethyl sulfoxide (DMSO) and used at 1 µM final concentration. Anti-PARP1 antibody (Cell Signaling Technology, Cat: 9542) was used at 1:5000. Anti-PARP2 antibody (Active Motif, Cat: 39044) was used at 1:2000. Anti-β-actin antibody (Sigma, Cat: A5441) was used at 1:20000. Anti-XRCC1 antibody (Novus Biologicals, Cat: NBP1-87154) was used at 1:5000. Anti-GFP antibody (Rockland, Cat: 600-301-215) was used at 1:1000. Anti-α-tubulin antibody (Sigma, Cat: CP06) was used at 1:1000. Anti-Pan-PAR binding reagent (Sigma, Cat: MABE1016) was used at 1:2000.

### Plasmids

The mRFP-C1-XRCC1 and pEGFP-C1-PARP2 plasmids were generously provided by Dr. Li Lan, currently at Duke University (31), and Dr. Xiaochun Yu, currently at Westlake University (32). The pEGFP-C1-PARP2-E545A and -R140A plasmids were described in our previous study (10). The plasmids encoding truncated PARP2 or XRCC1 were generated via PCR-mediated mutagenesis and subcloned into the pEGFP-C1 and mRFP-C1 vectors (Clontech), respectively. The protein sequence of the flexible GS-XTEN linker that was used to replace the linkers in XRCC1 is SGSET PGTSE SATPE SGGSG SSGGS GSSGG (N->C). All plasmids with targeted mutations were validated via Sanger sequencing.

### Western blot

RPE-1 cells and iMEFs were lysed with SDS lysis buffer (1% SDS, 10 mM HEPEs (pH=7.0), 2 mM MgCl_2_) supplemented with 0.2% Benzonase (Millipore, Cat: 71205) at room temperature for 10 min. To measure the auto-PARylation ability of different PARP2 variants, HEK 293T cells were transfected with plasmids encoding one of several GFP-tagged PARP2 variants. At 24 hours after transfection, the cells were lysed with NP-40 lysis buffer (25 mM Tris-HCl pH 8.0, 100 mM NaCl, 1% NP-40, 5 mM MgCl_2_, 10% Glycerol) supplemented with protease inhibitor cocktail (Roche, Cat: 11697498001), phosphatase inhibitors (Roche, Cat: 4906845001), 1 µM olaparib (Selleckchem, Cat: S1060), 1 µM PARG inhibitor PDD00017273 (Selleckchem, Cat: S8862) and 0.2% Benzonase at 4 °C for 30 minutes. Then, lysates were centrifuged at 4 °C, 13000 rpm for 15 minutes. The supernatants were collected, and protein concentrations were measured with the DC Protein Assay Kit II (BioRad, Cat: 5000112). The supernatants were mixed with SDS-PAGE loading buffer and incubated at 95 °C for 7 minutes before loaded on the gel. The 8% ∼ 12% gradient SDS-PAGE gel separated PARylated proteins of different sizes. After transfer via the standard protocol, the membranes were subjected to Western Blotting with indicated antibodies.

### Live-cell Imaging data collection and processing

Quantitative live-cell imaging analyses were performed as described previously (33) with minor modifications. Approximately 5 × 10^4^ RPE-1 cells or 1 × 10^5^ iMEFs were seeded onto a 35 mm glass-bottom plate. On the following day, the cells were transfected with plasmids encoding fluorescence-tagged PARP2 or XRCC1 using Lipofectamine 2000 (Invitrogen, Cat. 11668019) or Lonza 4D-Nucleofector™ X, following the manufacturer’s instructions. Live-cell imaging was conducted at 24 hours after initial transfection using a Nikon Ti Eclipse inverted microscope (Nikon Inc, Tokyo, Japan) equipped with the A1 RMP confocal microscope system and Lu-N3 Laser Units (Nikon Inc.). Only cells expressing the GFP- or RFP-tagged protein within a moderate yet reliable range (∼200–1000 a.u.) were selected for imaging. Laser micro-irradiation with a 405 nm laser at approximately 500 uW (affected area = 0.8 µm in diameter) and time-lapse imaging were performed using the NIS Element High Content Analysis software (Nikon Inc.) . Images were captured every 10 seconds after micro-irradiation for 5 minutes. The relative intensity at damaged sites was calculated as the ratio of the mean intensity at each micro-irradiated site to the corresponding mean intensity of the nucleus as background. Quantitative analysis was conducted using Fiji software, with at least eight individual cells per time point in each experiment. At least two independent experiments, each with eight cells, were conducted.

## RESULTS

### PARP1 and its catalytic activities promote the enrichment of PARP2 at the DNA damage foci

To study the kinetics of PARP2 at DNA damage sites, we established a live-cell imaging assay to monitor GFP-tagged PARP2 foci intensity following a 405nm laser micro-irradiation (10). The 405nm laser generates a mixture of predominantly single-stranded DNA breaks (SSBs) and a minor fraction of double-strand breaks (DSBs) measured by limited recruitment of GFP-KU, a DSB specific end-binding protein (34). While PARP1 and PARP2 both can be effectively activated by model DNA substrates *in vitro*, in cells, PARP1 is much more abundant and responsible for most damage-induced PARylation (13). Thus, to study the kinetics of GFP-PARP2 without the potentially confounding effect of endogenous PARP2 and PARP1, we examined GFP-tagged PARP2 foci formation in both PARP1 proficient (*PARP2 KO*) and deficient (*PARP1/2 DKO*) cells generated via CRISPR KO (10). We noted that micro-irradiation-induced GFP-PARP2 foci were significantly weaker in *PARP1/2 DKO* cells than in *PARP2 KO* RPE1 cells (Fig. 1A-C), suggesting that PARP1 promotes PARP2 foci formation. Moreover, in the absence of PARP1, PARP2 foci took longer to reach their maximum intensity (from 40 to 90 seconds, p<0.0001) (Fig. 1D), suggesting that PARP1 promotes early recruitment of PARP2. These cell lines have endogenous XRCC1. We also introduced RFP-XRCC1 as an indicator for the PAR levels (Fig. 1A). Consistent with PARP1 as a major responder, XRCC1 foci are much weaker in *PARP1/2 DKO* cells than in *PARP 2KO* cells (Fig. 1E). Next, we sought to understand whether the PARP1 protein or its activity, and by extension PAR, facilitates the recruitment of PARP2. Structural analyses identified an essential role of the Arginine-140 (R140) residue in the PARP2 WGR domain for DNA lesion binding (26). Our recent study showed that PARP2-R140A (RA) cannot bind DNA lesions and can only be recruited to the microirradiation site via the PARP1-dependent mechanism (10). Taking advantage of this mutation, we tested how PARP inhibition, which traps PARP1 protein while blocking PAR formation, impacts PAPR2-RA foci formation. FDA-approved PARPi, niraparib, successfully abolished DNA-damage induced PAR formation measured by reduced XRCC1 foci and markedly decreased PARP2-RA foci (Fig. 1F-I), suggesting PARP1 activity, and by extension PAR, promotes PARP2 foci formation. We considered two possible mechanisms: direct binding to PAR or indirect recruitment through one or several PAR-binding mediators.

**Figure 1.**
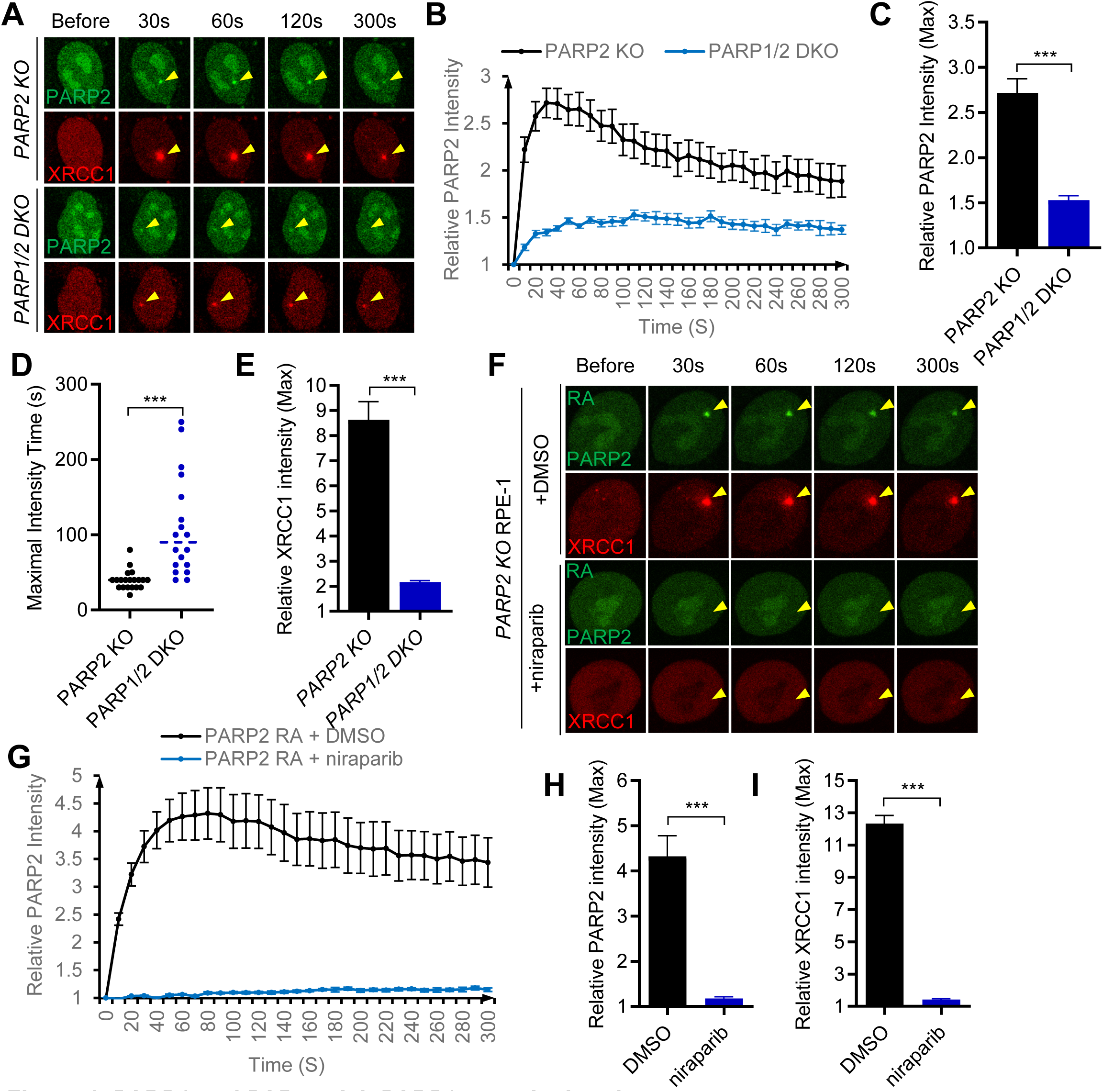
PARP1 and PAR enrich PARP2 near the breaks. (A) The representative live-cell images of laser-induced GFP-PARP2 and RFP-XRCC1 foci in *PARP2 KO* or *PARP1/2 DKO* RPE-1 cells. The yellow arrowhead points to the areas of micro-irradiation. (B) The relative intensity kinetics of GFP-PARP2 in *PARP2 KO* or *PARP1/2 DKO* RPE-1 cells. (C) The maximal relative intensity of GFP-PARP2. (D) The time it took for the micro irradiation-induced GFP-PARP2 foci to reach maximal intensity was plotted from images collected from 20 *PARP2 KO* cells and 18 *PARP1/2 DKO* cells. Mann-Whitney test was used to calculate p-values. (E) The maximal relative intensity of RFP-XRCC1. (F) Representative live-cell images of laser-induced GFP-PARP2^RA^ and RFP-XRCC1 foci in *PARP2 KO* RPE-1 cells in the presence of 1µM niraparib for 1h. The yellow arrowheads indicate the areas of micro-irradiation. (G) The relative intensity kinetics of GFP-PARP2^RA^ in *PARP2 KO* cells w/ or w/o niraparib. (H-I) The maximal relative intensity of (H) GFP-PARP2^RA^ and (I) RFP-XRCC1. The bars and error bars represent means and standard errors of mean (SEM) from one out of three representative experiments with n > 8 cells per experiment. The two-tailed unpaired student’s t-test was used to calculate the p-values. ***p<0.001.

### XRCC1 promotes the PARP1 and PAR-dependent enrichment of PARP2 to DNA damage sites

Upon DNA damage, XRCC1 is rapidly recruited to the sites of DNA damage via the direct interaction between its central BRCT1 domain and PAR (35). Given the prominent role of XRCC1 as a PAR responder, we compared micro-irradiation-induced GFP-PARP2 foci in *PARP1^+/+^PARP2^+/+^*(WT), *PARP1 ^KO^*, *XRCC1 ^KO^,* and *PARP1 and XRCC1 double KO* RPE-1 cells generated via CRISPR KO and validated via Western blotting (Fig. 2A). These cells have endogenous PARP2, and the GFP-PARP2 needs to compete with endogenous PARP2 for recruitment, thus forming weaker foci than in PARP2 KO cells (Fig. 1). Strikingly, the loss of XRCC1 markedly attenuated the micro-irradiation-induced GFP-PARP2 foci (Fig. 2B-D). This result was unexpected since XRCC1 deficiency is known to cause the accumulation of unrepaired DNA breaks, hyper-recruitment, and hyperactivation of PARP1 (21,22,36). Moreover, co-deletion of PARP1 and XRCC1 did not further diminish GFP-PARP2 foci, suggesting an epistatic model (Fig. 2B-D). Furthermore, the ectopic expression of RFP-XRCC1 in *XRCC1 KO* cells restored GFP-PARP2 RA foci formation at DNA damage sites (Fig. 2E-G). The DNA-binding deficient GFP-PARP2-RA (10) was used to avoid the confounding effects of DNA-dependent recruitment of PARP2. The importance of XRCC1 in promoting PARP2 recruitment is not limited to RPE-1/human cells and was also found in murine embryonic fibroblasts (MEFs), where the loss of Xrcc1 also significantly attenuated GFP-PARP2 foci formation (Fig. S1A-D). Together, these data identify XRCC1 as a key mediator for PAR-dependent enrichment of PARP2 at DNA damage sites.

**Figure 2.**
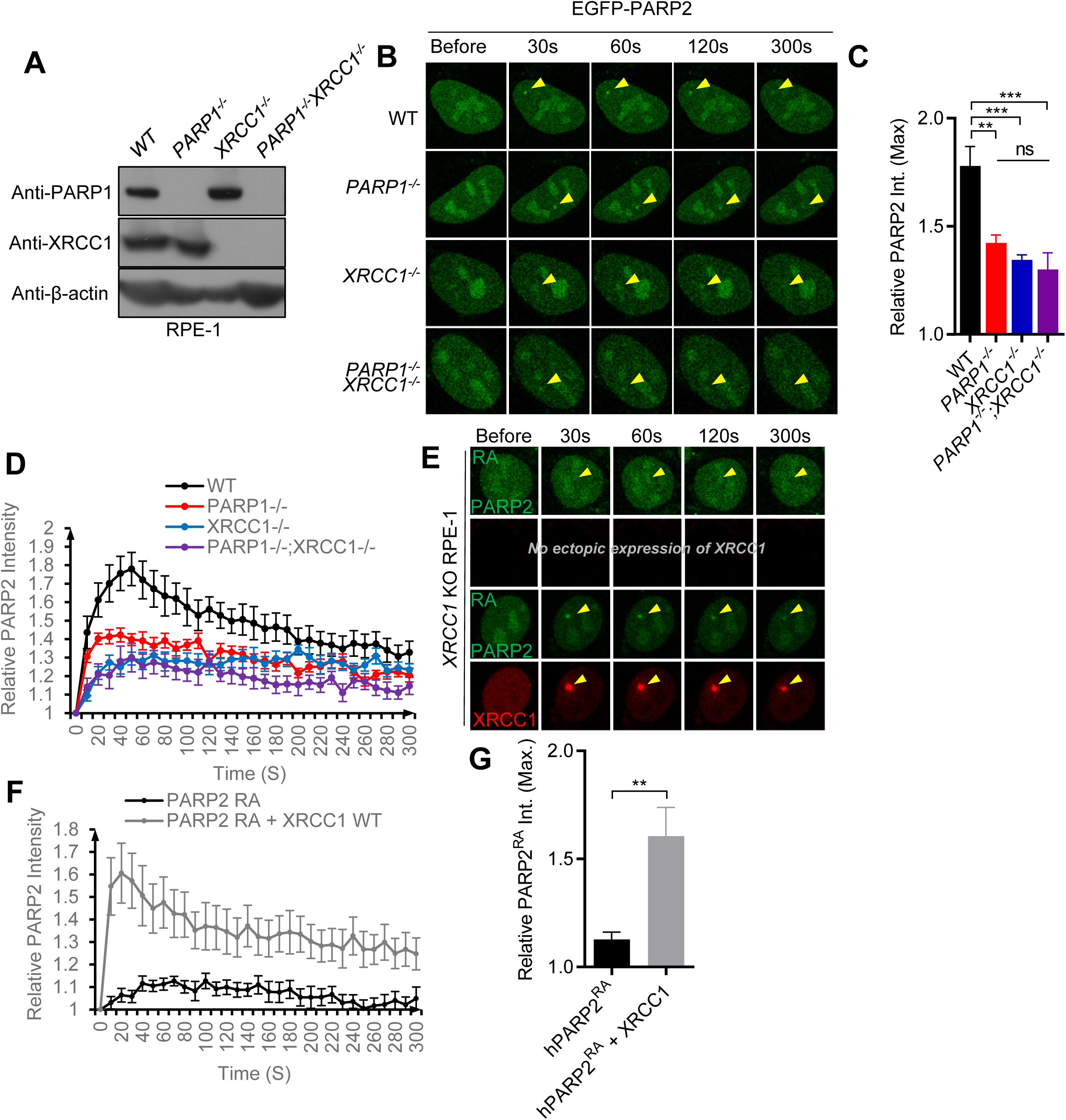
XRCC1 is required for PAR-dependent enrichment of PARP2 near the breaks. (A) Western blot analysis with anti-PARP1, anti-XRCC1, and anti-β-actin in *WT, PARP1 KO, XRCC1 KO, and PARP1/XRCC1 DKO* RPE-1 cells. (B) Representative live-cell images of laser-induced GFP-PARP2 foci in *WT, PARP1 KO, XRCC1 KO, and PARP1/XRCC1 DKO* RPE-1 cells. The yellow arrowhead points to the area of micro-irradiation. (C) The maximal relative intensity of GFP-PARP2. (D) The relative intensity kinetics of GFP-PARP2 at DNA damage sites in *WT, PARP1 KO, XRCC1 KO, and PARP1/XRCC1 DKO* RPE-1 cells. The points and error bars represent means and SEM from one out of two representative experiments with n > 8 cells per experiment. (E) Representative live-cell images of GFP-PARP2^RA^ and RFP-XRCC1 in *XRCC1 KO* RPE-1 cells. (F) The relative intensity kinetics of GFP-PARP2^RA^ at DNA damage sites. (G) The maximal relative intensity of GFP-PARP2^RA^ w/ and w/o RFP-XRCC1. The bars and error bars represent means and SEM from one out of three representative experiments with n > 8 cells per experiment. The two-tailed unpaired student’s t-test was used to calculate the p-values. ns, p> 0.05; **p<0.01; ***p<0.001.

### XRCC1-mediated recruitment of PARP2 requires the CAT domain of PARP2

To identify the domains of PARP2 that contribute to this XRCC1-mediated enrichment, we generated two truncated mutants, the PARP2^ΔWGR^ and PARP2^ΔCAT^, in which we removed either the WGR domain implicated in DNA binding or the C-terminal CAT domain, respectively (Fig 3A). A flexible GS-XTEN linker replaced the WGR domain in the PARP2^ΔWGR^ vector to minimize the impact on protein folding and stability (Fig 3A). To avoid the influence of endogenous PARP2, we compared the kinetics of PARP2^ΔWGR,^ PARP2^ΔCAT^, and full-length PARP2 (PARP2^WT^) in *PARP2 KO* cells with concurrent expression of RFP-XRCC1 that labels the damage sites. Overexpression of XRCC1 also augments the PAR and XRCC1-dependent recruitment of PARP2, providing a sensitive platform to determine the domain requirement of PARP2. In comparison to GFP-PARP2^WT^, GFP-PARP2^ΔWGR^ displayed slightly delayed kinetics and a moderately reduced (but not significant) intensity, likely reflecting the role of WGR in DNA damage-mediated recruitment and activation of PARP2 (Fig. 3B-D). Correspondingly, maximal XRCC1 foci intensity is slightly lower in cells expressing PARP2^ΔWGR^ than those expressing PARP2^WT^ (Fig. 3E). In contrast, GFP-PARP2^ΔCAT^ failed to form foci despite over-expression of XRCC1 and robust XRCC1 foci (Fig. 3B-E). Notably, both GFP-PARP2^ΔCAT^ and GFP-PARP2^ΔWGR^ contain the entire NTR, suggesting that NTR is neither necessary nor sufficient to support PARP2 recruitment, consistent with prior comparison of PARP2^WT^ and PARP2^ΔNTR^.(10) Next, we tested whether the PARP2-CAT domain lacking the DNA-binding motifs (PARP2^CAT^), is sufficient to support XRCC1-dependent recruitment of PARP2. The nuclear localization signal (NLS) of PARP2 resides in the NTR (6) and is missing in the PARP2^CAT^, explaining why GFP-PARP2^CAT^ was found in both the nucleus and cytoplasm (Fig. 2F). In the absence of XRCC1, GFP-PARP2^CAT^ did not form measurable foci (Fig. 3F-H), consistent with lack-of-DNA binding. In cells with reconstitution of RFP-XRCC1, GFP-PARP2^CAT^ form robust and DNA-damage-induced foci (p<0.001) (Fig. 3F-H). Together, the results show that the PARP2-CAT domain is necessary and sufficient for the PAR and XRCC1-dependent recruitment of PARP2 to the vicinity of DNA damage sites.

**Figure 3.**
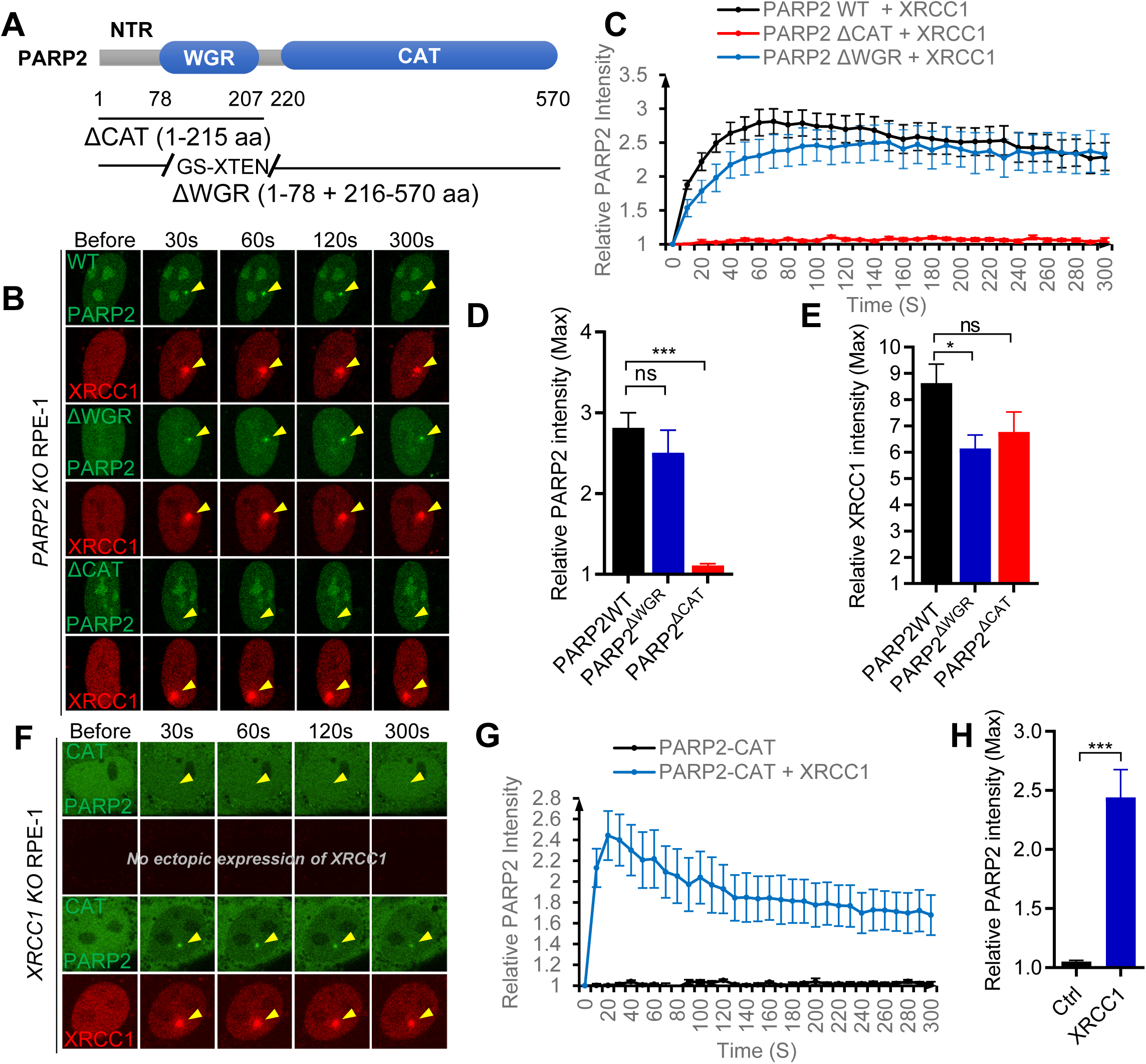
XRCC1-dependent enrichment of PARP2 requires the CAT domain of PARP2. (A) The schematic of PARP2 domains and truncated PARP2 with aa marked. (B) Representative live-cell images of laser-induced GFP-PARP2 and RFP-XRCC1, GFP-PARP2^ΔWGR^ and RFP-XRCC1, and GFP-PARP2^ΔCAT^ and RFP-XRCC1 foci in *PARP2 KO* RPE-1 cells. (C) The relative intensity kinetics of GFP-PARP2, GFP-PARP2^ΔWGR^, and GFP-PARP2^ΔCAT^ at DNA damage sites. (D) The maximal relative intensity of GFP-PARP2, GFP-PARP2^ΔWGR^, and GFP-PARP2^ΔCAT^. (E) The maximal relative intensity of RFP-XRCC1. (F) Representative live-cell images of GFP-PARP2^CAT^ alone and GFP-PARP2^CAT^ and RFP-XRCC1 in *XRCC1 KO* RPE-1 cells. (G) The relative intensity kinetics and (H) the maximal relative intensity of GFP-PARP2^CAT^ w/ and w/o RFP-XRCC1. The bars and error bars represent means and SEM from one out of three representative experiments with n > 8 cells per experiment. The two-tailed unpaired student’s t-test was used to calculate the p-values. ns, p> 0.05; *p<0.05; ***p<0.001.

### XRCC1-mediated recruitment of PARP2 does not require PARP2 catalytic activity

XRCC1 binds to PAR via its center BRCT1 domain to recruit itself and its cargos to the DNA damage sites(35). Given one important function of the PARP2 CAT domain is to synthesize the PAR chain, we tested whether PARP2’s catalytic activity and auto-PARylation are required for the XRCC1-mediated enrichment of PARP2 at DNA damage sites. We took advantage of the previously characterized catalytically inactive PARP2^E545A^, which lacks the PARylation-specific E545 in the H-Y-E catalytic triad (10). While both PARP2^WT^ and PARP2^E545A^ formed DNA damage-induced foci, GFP-PARP2^WT^, but not GFP-PARP2^E545A^ supported XRCC1 foci formation in *PARP1/2 DKO* cells, validating the loss of PARylation activity in the PARP2^E545A^ (Fig. S2A-E). Moreover, in comparison with PARP2^WT^, both PARP2^CAT^ and PARP2^E545A^ were not auto-PARylated when ectopically expressed in HEK239T cells with endogenous PARP1 and PARP2 (Fig. 4A). This observation also supports the strict intramolecular auto-PARylation of PARP2, rather than intermolecular auto-PARylation (Fig. 4A). Nevertheless, GFP-PARP2^E545A^ forms XRCC1-dependent and DNA damage-induced foci (Fig. 4B-D) (p<0.001, student’s t-test). Thus, XRCC1 promotes PAR-dependent enrichment of PARP2 at DNA damage sites independent of PARP2 catalytic activity or PARP2 auto-PARylation, suggesting a role of XRCC1 in enriching PARP2 before its activation

**Figure 4.**
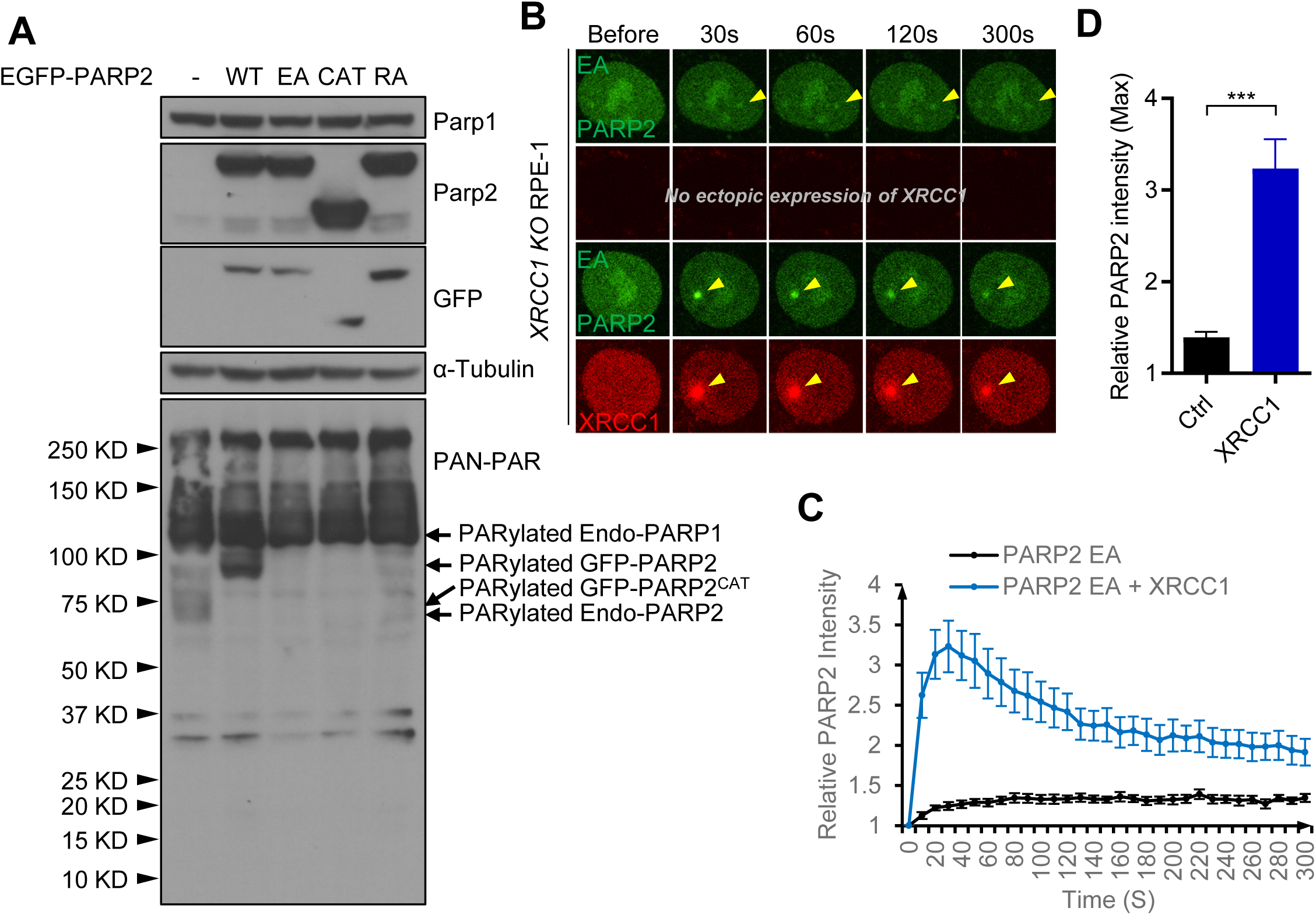
XRCC1-dependent enrichment of PARP2 near the breaks does not require PARP2 catalytic activity. (A) Western blot to measure the PARylation level of GFP-PARP2^WT^, GFP-PARP2^E545A^, GFP-PARP2^CAT^ and GFP-PARP2^R140A^ in wild-type 293T cells (B) Representative live-cell images of laser-induced GFP-PARP2^E545A^ alone, and GFP-PARP2^E545A^ and RFP-XRCC1 foci in *XRCC1 KO* RPE-1 cells. (C) The relative intensity kinetics and (D) the maximal relative intensity of GFP-PARP2^E545A^ w/ or w/o RFP-XRCC1 at DNA damage sites. The bars and error bars represent the means and SEM from one out of two independent experiments with n > 8 cells per experiment. The two-tailed unpaired student’s t-test was used to calculate the p-values. ***p<0.001.

### The BRCT1 domain of XRCC1 and its PAR binding pocket are necessary but insufficient for the recruitment of PARP2

The XRCC1 protein contains the N-terminal domain (NTD) and two BRCT domains connected with two flexible linkers (Fig. 5A). To identify the domains of XRCC1 necessary for PARP2 recruitment, we generated a series of plasmids expressing truncated RFP-tagged XRCC1. The BRCT1 domain of XRCC1 has been found to bind to both PARP1 and PARP2 (37) and is required for PAR binding (35). While the RFP-XRCC1^BRCT1^ (aa 161-406, BRCT1 fragment) in *XRCC1 KO* RPE-1 cells formed robust DNA damage-induced foci indistinguishable from RFP-XRCC1^WT^, it failed to recruit GFP-PARP2 to the foci (Fig. 5B-E). The XRCC1^BRCT1^ binds PAR via its phosphorylation binding pocket. Previous analyses showed that alanine substitution at Arginine 335 and Lysine 369 within the BRCT1 domain abolishes PAR binding in the context of full-length XRCC1 (38). RFP-tagged XRCC1^RK^ (R335A/K369A) was not able to form DNA damage-induced foci likeRFP-XRCC1^WT^ or RFP-XRCC1^BRCT1^ (Fig. 5E) and could not support GFP-PARP2 foci formation in *XRCC1 KO* cells (Fig. 5B-D). Thus, the BRCT1 domain of XRCC1 is necessary but insufficient to support PAR-dependent recruitment of PARP2. The results support a working model in which XRCC1 binds to PAR through its BRCT1 domain, and then another region of XRCC1 binds to PARP2 and recruits PARP2 to the breaks.

**Figure 5.**
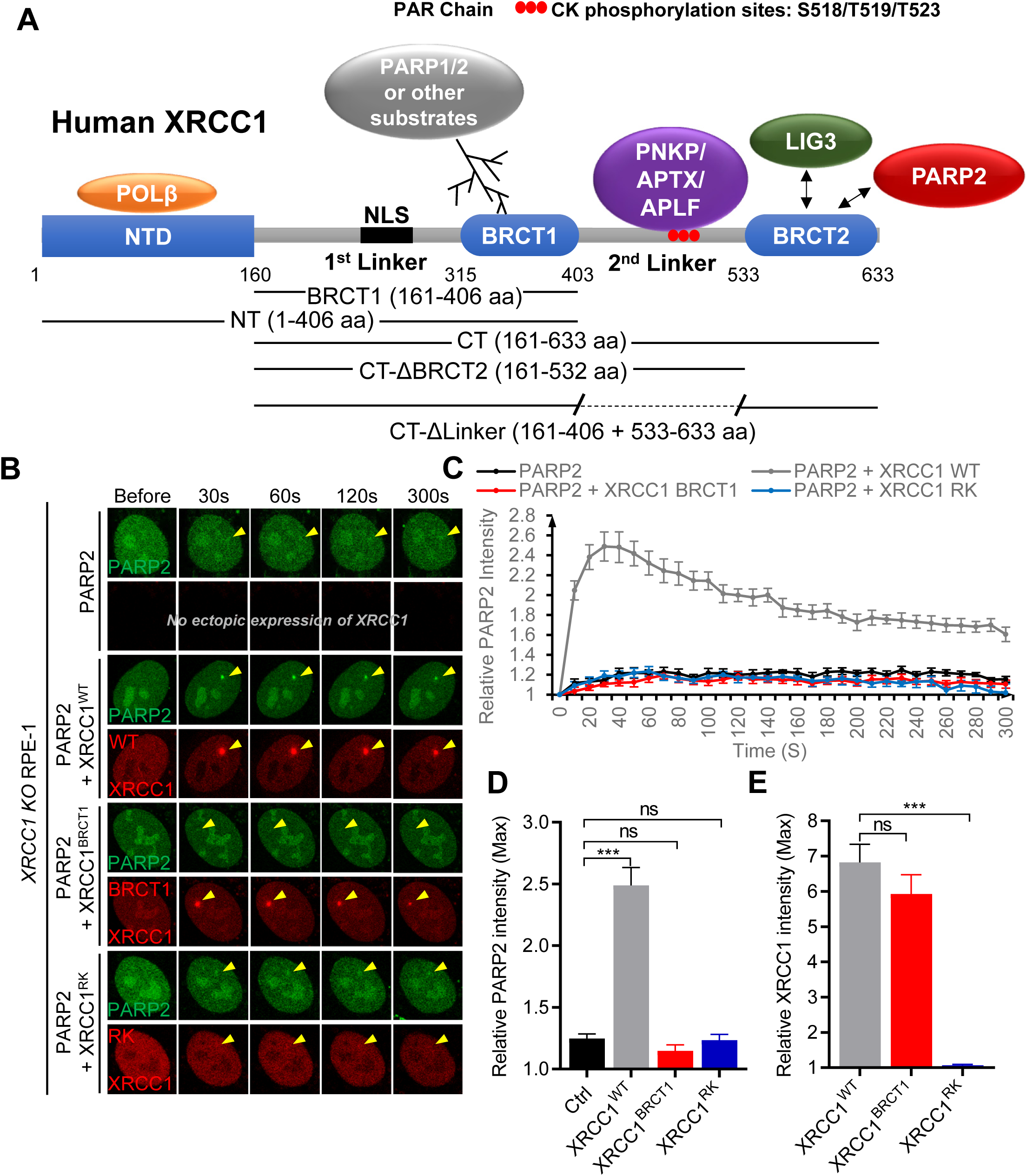
The BRCT1 domain of XRCC1 is essential but not sufficient for PAR-dependent recruitment of PARP2. (A) The schematic of XRCC1 domains and their interaction partners PARP1/2, PolB, Lig3, PNKP, APTX, and APLF with aa marked for human XRCC1. The sequence information of truncated XRCC1 mutants is labeled. (B) Representative live cell images of laser-induced GFP-PARP2 alone, GFP-PARP2 and RFP-XRCC1, GFP-PARP2 and RFP-XRCC1^BRCT1^, GFP-PARP2 and RFP-XRCC1^RK^ (R335A/K369A) foci in XRCC1 *KO* RPE-1 cells. The yellow arrowhead points to the areas of micro-irradiation. (C) The relative intensity kinetics of GFP-PARP2 alone or with RFP-XRCC1, RFP-XRCC1^BRCT1^, or RFP-XRCC1^RK^ at DNA damage sites. (D) The maximal relative intensity of GFP-PARP2 and (E) RFP-XRCC1, RFP-XRCC1^BRCT1^, and RFP-XRCC1^RK^. The bars and error bars represent the means and SEM from one of three independent experiments with n > 8 cells per experiment. The two-tailed unpaired student’s t-test was used to calculate the p-values. ns, p>0.05; ***p<0.001.

### The BRCT2 domain of XRCC1 mediates the recruitment of PARP2

Next, we tested whether the N- or the C-terminal region of XRCC1 can recruit PARP2 by generating an N-terminal truncated (XRCC1^CT^, a.a. 161-633) and a C-terminal truncated (XRCC1^NT^, aa 1-406) XRCC1. Both RFP-XRCC1^CT^ and RFP-XRCC1^NT^ contain the BRCT1 domain and the endogenous nuclear localization signal (Fig. 5A) and form robust DNA damage-induced foci (Fig. 6A,D). But only the RFP-XRCC1^CT^, not the RFP-XRCC1^NT^ supported the damage-induced PARP2 foci, suggesting the C-terminus of XRCC1, including BRCT1, 2^nd^ linker, and BRCT2, is sufficient to recruit PARP2 (Fig. 6A-C). The 2^nd^ linker contains a cluster of CK2 phosphorylation sites that can be recognized by PNKP, APTX, and APLF (39) (Fig. 5A). To determine whether the 2^nd^ linker, the BRCT2 domain, or both are required for PARP2 recruitment, we removed the BRCT2 (XRCC1^CT-ΔBRCT2^) or substituted the 2^nd^ linker with a GS-XTEN flexible linker (XRCC1 ^CT-ΔLinker^) (Fig. 5A). RFP-XRCC1 ^CT-ΔBRCT2^ formed bright and robust foci but could not support the recruitment of PARP2 (Fig. 6E-H), suggesting a role of BRCT2 in PARP2 recruitment. Meanwhile, the loss of the 2^nd^ linker (XRCC1 ^CT-ΔLinker^) compromised both the DNA damage-induced XRCC1 foci and PARP2 foci, suggesting that the 2^nd^ linker contributed to the stabilization of the XRCC1 foci (Fig. 6E-H). While the results established a critical role of the XRCC1 BRCT2 domain in PARP2 recruitment, whether the XRCC1 linker directly contributes to PARP2 recruitment cannot be unequivocally determined.

**Figure 6.**
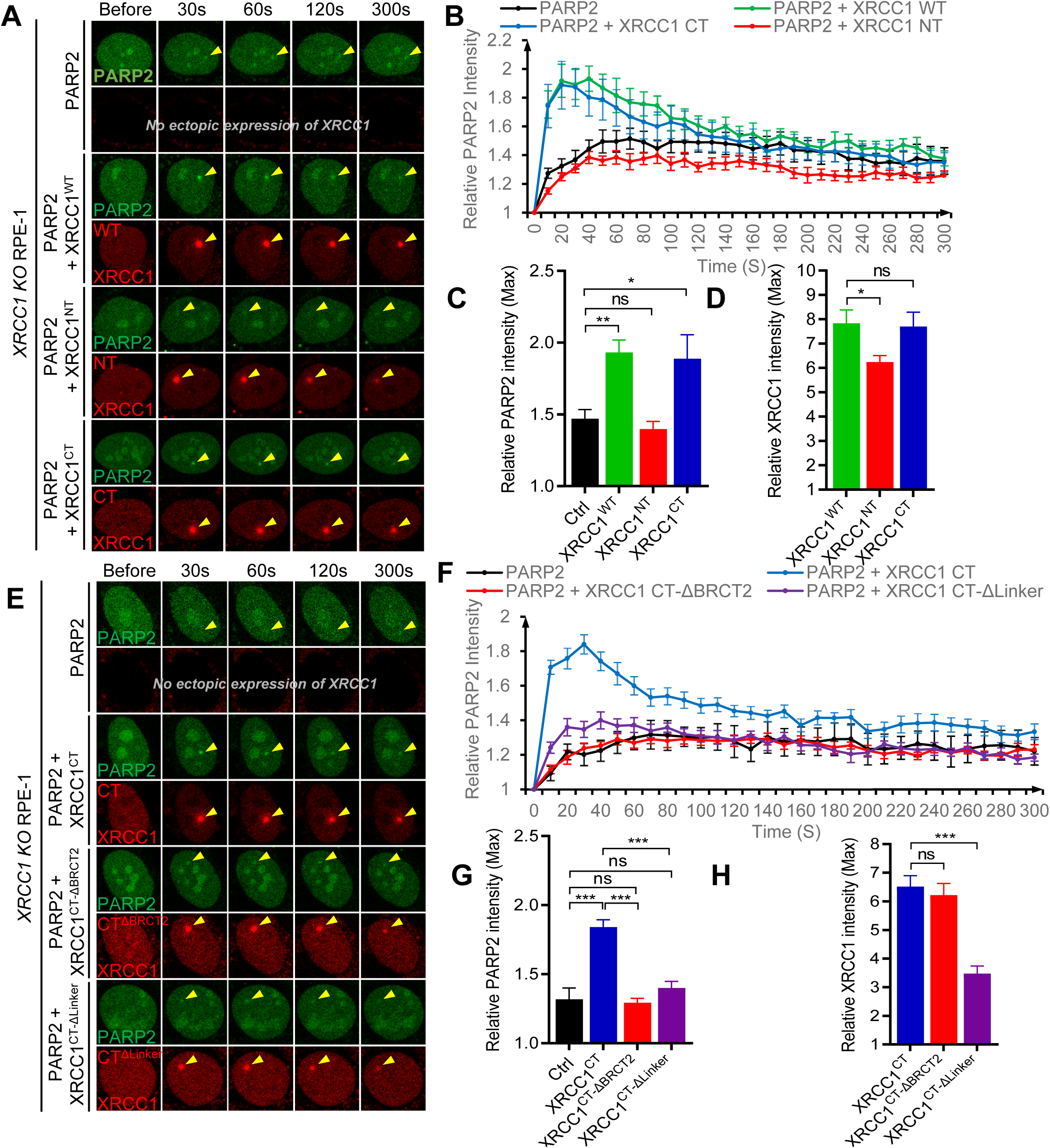
The XRCC1 BRCT2 domain interacts with and recruits PARP2. (A) Representative live-cell images of laser-induced GFP-PARP2 alone, GFP-PARP2 and RFP-XRCC1^WT^, GFP-PARP2 and RFP-XRCC1^NT^, GFP-PARP2 and RFP-XRCC1^CT^ foci in *XRCC1 KO* RPE-1 cells. The yellow arrowheads point to the areas of micro-irradiation. (B) The relative intensity kinetics of GFP-PARP2 alone or with RFP-XRCC1, RFP-XRCC1^NT^ or RFP-XRCC1^CT^ at DNA damage sites. (C) The maximal relative intensity of GFP-PARP2 and (D) RFP-XRCC1, RFP-XRCC1^NT^, and RFP-XRCC1^CT^. (E) Representative images of GFP-PARP2, GFP-PARP2 and RFP-XRCC1^CT^, GFP-PARP2 and XRCC1^CT-ΔBRCT2^, and GFP-PARP2 and RFP-XRCC1^CT-ΔLinker^ foci in *XRCC1 KO* RPE-1 cells. (F) The relative intensity kinetics of GFP-PARP2 alone or with RFP-XRCC1^CT^, RFP-XRCC1^CT-ΔBRCT2^, or RFP-XRCC1^CT-ΔLinker^. (G) The maximal intensity of GFP-PARP2 and (H) RFP-XRCC1^CT^, RFP-XRCC1^CT-ΔBRCT2^, or RFP-XRCC1^CT-ΔLinker^. The bars and error bars represent the means and SEM from one of three independent experiments with n > 8 cells per experiment. The two-tailed unpaired student’s t-test was used to calculate the p-values. ns, p> 0.05; *p<0.05; **p<0.01; ***p<0.001.

### XRCC1 BRCT2 domain directly recruits PARP2 independent of LIG3

The BRCT2 of XRCC1 is known to bind to LIG3. Specifically, the arginine 564 residue within the BRCT2 domain is required for XRCC1-LIG3 heterodimer formation *in vitro* (40). To determine whether LIG3 is required as a mediator for XRCC1-dependent recruitment of PARP2, we introduced the R564A mutation into full-length RFP-XRCC1 (XRCC1^R564A^). To our surprise, in *XRCC1 KO* cells, ectopically expressed RFP-XRCC1^R564A^ formed robust DNA damage-induced foci and fully retained damage-induced LIG3 foci (Fig. S3A-D), but failed to recruit PARP2 to the DNA damage sites (Fig. 7A-D). In this context, XRCC1 also interacts with LIG3 on several additional residues, including Arg560 and Leu539 (40), which might contribute to the residual LIG3 foci in the overexpression system. However, in *XRCC1 KO* cells reconstituted with XRCC1^R564A^, PARP2 foci were not fully restored, suggesting that the R564 in the BRCT2 domain of XRCC1 is essential for XRCC1-dependent recruitment of PARP2. This interaction is independent of PARP2 catalytic activity and is mediated by a direct interaction between XRCC1 and PARP2.

**Figure 7.**
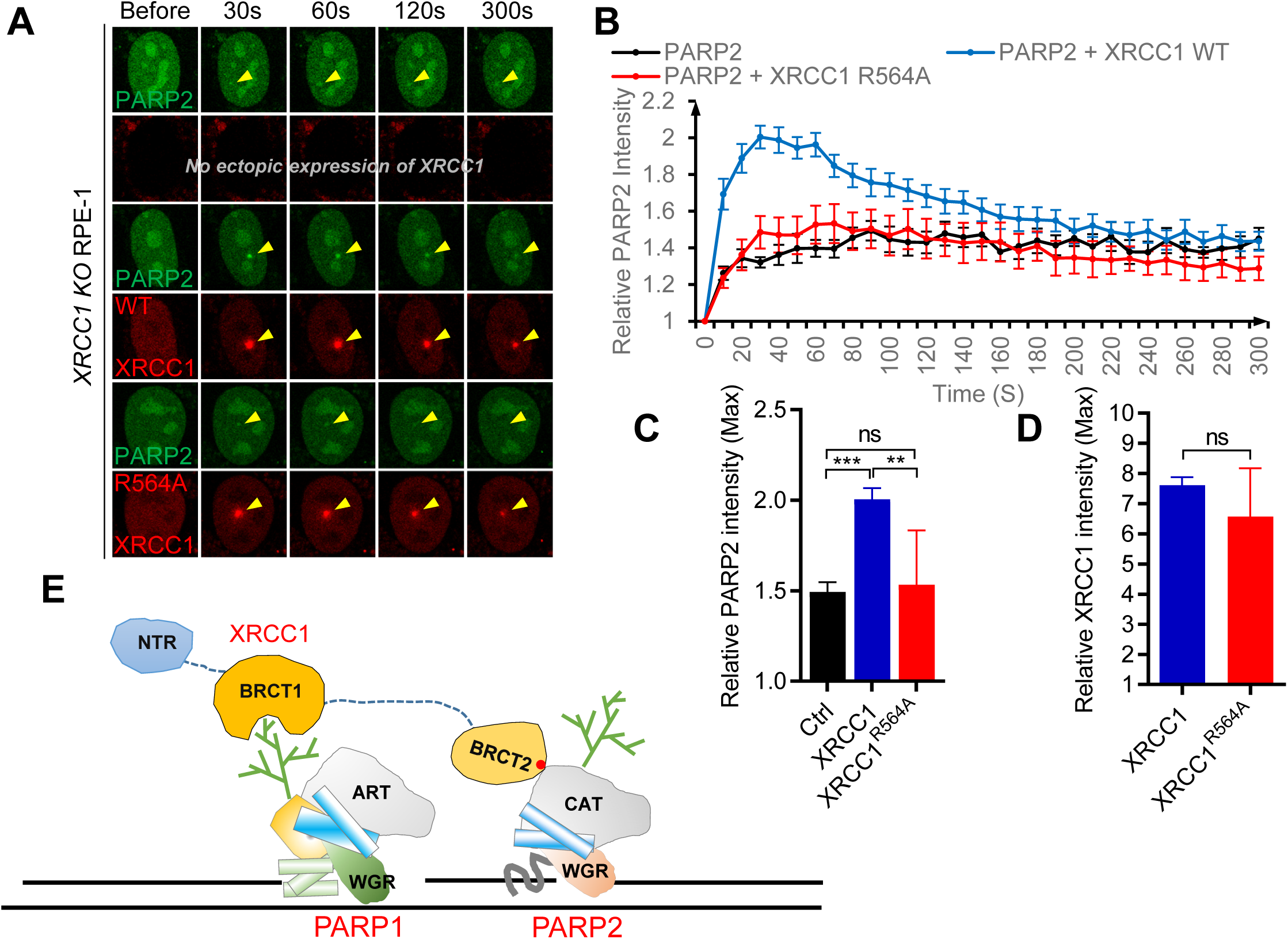
The phosphorylation binding moiety on the XRCC1 BRCT2 domain is necessary for PARP2 enrichment. (A) Representative live-cell images of laser-induced foci of GFP-PARP2 alone, GFP-PARP2 and RFP-XRCC1, and GFP-PARP2 and RFP-XRCC1^R564A^ in *XRCC1 KO* RPE-1 cells. The yellow arrowheads mark the areas of micro-irradiation. (B) The relative intensity kinetics of GFP-PARP2 alone or with RFP-XRCC1, or RFP-XRCC1^R564A^. (C) The maximal relative intensity of GFP-PARP2 and (D) RFP-XRCC1 and RFP-XRCC1^R564A^. The bars and error bars represent means and SEM from one out of three representative experiments with n > 8 cells per experiment. The two-tailed unpaired student’s t-test was used to calculate the p-values. ns, p> 0.05; **p<0.01; ***p<0.001. (E) A working model shows that XRCC1 binds to PAR via BRCT1 and uses BRCT2 to recruit PARP2.

## DISCUSSION

Based on sequence homology, PARP2 was cloned 25 years ago as the enzyme responsible for the residual PARylation activity in PARP1-deficient cells (13,41). Subsequently, genetic data solidified the role of PARP2 as a “backup” for PARP1 in its absence (11,12). However, several differences between PARP1 and PARP2 have been noted recently. With its smaller and simpler DNA binding motif, PARP2 is selectively activated by 5’ phosphorylated nicks, unlike the diverse DNA lesions (*e.g.*, DSBs, gaps, etc) that activate PARP1 (7). Correspondingly, PARP2 also has a faster exchange rate, as measured by fluorescence recovery after photobleaching (9,10). Moreover, PARP2 is less robust in its activity and much less abundant (42). While all of these observations support a backup and potentially much-limited role of PARP2, PARP2 has been found to contribute to many DNA repair pathways in which PARP1 is also involved, including base excision repair (BER), non-homologous end joining (NHEJ), and the recruitment and activation of PNKP and others (28,43,44). Moreover, deficiency of PARP2, but not PARP1, compromises erythropoiesis, T lymphocyte activation, and spermatogenesis *in vivo* (45–47). How could PARP2 access DNA lesions in the presence of abundant PARP1? Here, we propose that in addition to direct DNA binding with its WGR domain(6,10,26), PARP2 is also enriched at the vicinity of the breaks via a PARP1- and PAR-dependent mechanism (10,32). Specifically, we found that XRCC1 plays an essential role in the PARP1- and PAR-dependent enrichment of PARP2 by simultaneously binding to PAR via its central BRCT1 domain and PARP2 via its C-terminal BRCT2 domain. This bridging function of XRCC1 requires the phos-binding pocket in the BRCT1 domain and the CAT domain of PARP2 but is independent of PARP2 activity or auto-PARylations (Fig. 4B-D). Based on these data, we propose a working model (Fig. 7E): PARP1 is immediately recruited and activated upon DNA breaks. The resulting PAR chains recruit XRCC1, which, in addition to bringing other SSBR proteins, also enriches PARP2 near the breaks, where PARP2 can be activated to generate more PAR and forms a positive feedback loop until the lesion is repaired. This model might explain how the CAT domain could promote WGR-mediated DNA binding and foci formation by PARP2 despite lacking DNA binding ability (48). XRCC1-deficiency leads to neurological diseases in both human and mouse models (47–49), partly through hyperactivation of PARP1. This model also explains why XRCC1 deficiency is more devastating than PARP1 deficiency and why XRCC1 deficiency does not cause hyperactivation of PARP2. The recruitment and activation of PARP2 through XRCC1 might compete with PARP1 for DNA binding, promoting PARP1 release, in addition to faciliating repair. This might also contribute to the therapeutic and toxic effect of PARPi, which would also attenuate XRCC1-mediated recruitment of PARP2. Upon DNA damage, XRCC1 binds to PAR chains and acts as a scaffold to bring a series of SSBR enzymes to support end-modification, gap-filling, and ligation. Here, we show that XRCC1 is also a PAR-signal amplifier by recruiting PARP2 to the vicinity of the breaks. Given the ability for XRCC1 to recruit inactive PARP2, we propose that XRCC1-mediated enrichment of PARP2 occurs before PARP2 activation. Conveniently, the XRCC1-binding and DNA-binding of PARP2 are mediated by different domains of PARP2. As such, XRCC1 increases the local concentration of PARP2 near DNA breaks, allowing a smooth transition to DNA binding and to the promotion of PARP2 activation. Correspondingly, PARP1, PAR (blocked by PARPi), and XRCC1 all contribute to the early recruitment of PARP2 (Fig. 2A-C) (10). By doing so, through PARP2, XRCC1 might increase PAR chains’ complexity and extend PAR signaling’s duration. Temporally, PARP2 preferentially binds to 5’ phosphorylated nicks, the substrate of DNA ligase at the very last step of SSBR (7,44,49). Both LIG3 and PARP2 bind to the BRCT2 domain of XRCC1, likely not simultaneously. Further research is needed to understand how XRCC1 spatially and temporally coordinates SSBR step by step. Regarding complexity, Qian Chen *et al.* reported that PARP2 preferentially generates a branched PAR chain, which recruits APLF via its tandem PAR-binding zinc finger (PBZ) domains (32). Notably, APLF also binds to the CK2-phosphorylated motif adjacent to the XRCC1’s BRCT2 domain (35,50), suggesting that XRCC1 might promote APLF through at least two different mechanisms. This finding might also explain why the 2^nd^ linker of XRCC1 stabilizes XRCC1 at the breaks (Fig. 6E-G).

Together, these findings unveil a new function of XRCC1 in augmenting PARP2 recruitment in response to PARP1 activation and explain why PARP1, but not PARP2, is aggregated and hyperactivated in XRCC1-deficient cells, with implications in the development of PARP inhibitors and the pathological changes in XRCC1-deficiency.

## DATA AVAILABILITY

Supplementary Data will be available online.

## AUTHOR CONTRIBUTIONS

X.L. and S.Z conceptualized the initial project. XL and K.L. designed and. XL, KL, and K.W. performed the experiments, data analysis, and validation, and they prepared the first draft. B.J.L supports the projects with reagants and cell line inquiry and maintenance. S.Z. X.L and B.L. edited the drafts with inputs from all co-authors.

## ACKNOWLEDGEMENTS

We thank Drs. Theresa Swayne and Haojie Ji for their advice on confocal microscopy, Drs. Li Lan, Xiaochun Yu, and Keith Caldecott for generously providing cell lines, plasmids, and protocols, and the technical help from other Zha lab members. Due to space limitations, we could not cite all the original publications and reviews when necessary.

## FUNDING

This work was supported by the National Institutes of Health CA271595, CA 226852, and CA293675 to S.Z. This research used the co-focal microscopy facility (for imaging acquisition) and transgenic mouse facility (to generate immortalized MEFs of various genotypes), partly funded through the NIH/NCI Cancer Center Support Grant P30CA013696 to Herbert Irving Comprehensive Cancer Center (HICCC) of Columbia University.

## Supplementary Figure Legend

**Figure S1.**
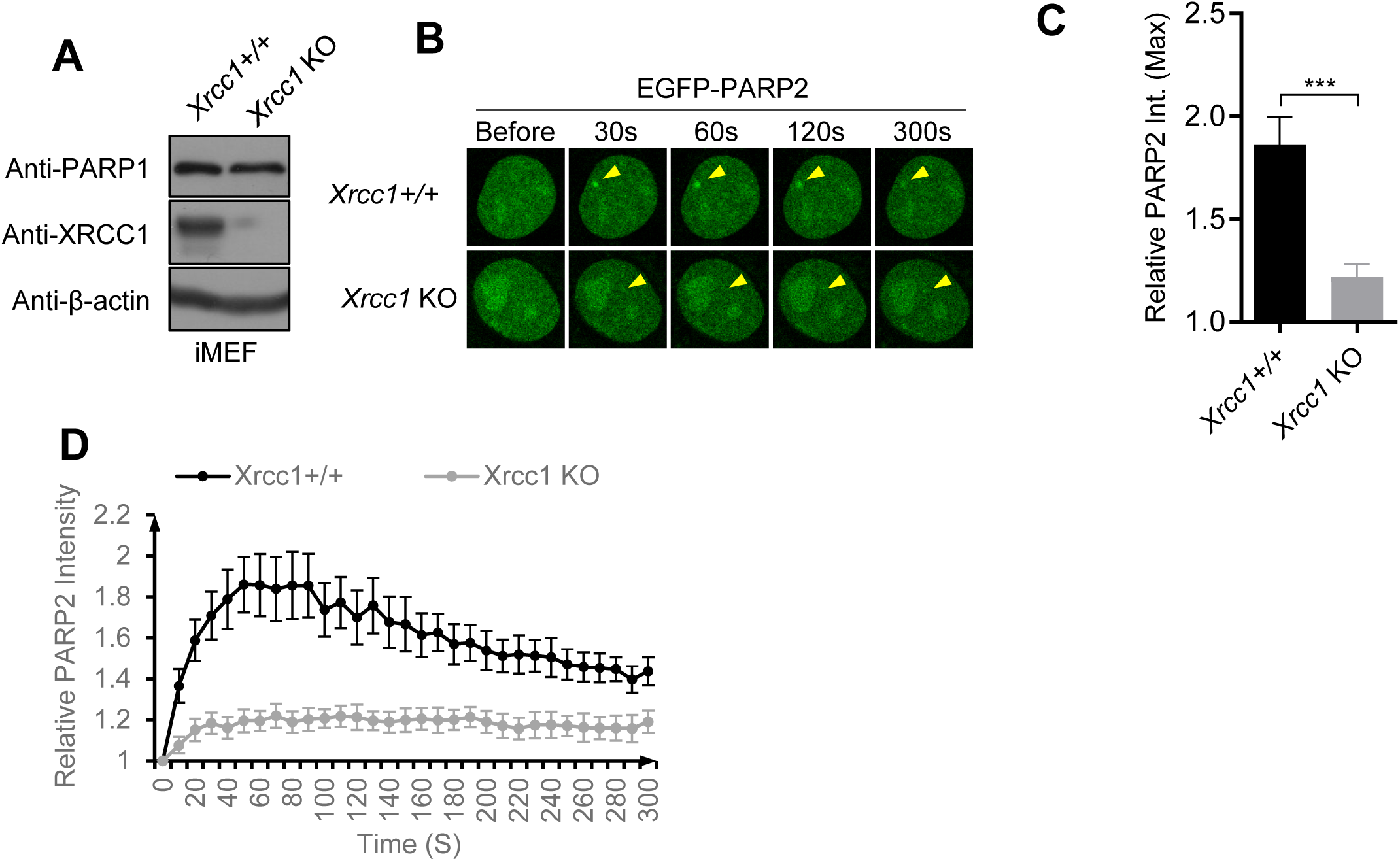
XRCC1 mediates PARP2 recruitment to DNA damage sites in iMEF. (A) Western blot analyses with anti-PARP1, anti-XRCC1, and anti-β-actin antibodies on *Xrcc1^+/+^* and *Xrcc1^-/-^* iMEFs. (B) Representative live-cell images of laser-induced GFP-PARP2 foci in *Xrcc1+/+* and *Xrcc1 KO* iMEFs. The yellow arrowheads point to the area of micro-irradiation. (C) The maximal relative intensity and (D) the relative intensity kinetics of GFP-PARP2 at DNA damage sites in *Xrcc1+/+* and *Xrcc1 KO* iMEFs. The bars and error bars represent means and SEM from one out of two representative experiments with n > 8 cells per experiment. The two-tailed unpaired student’s t-test was used to calculate the p-values. ***p<0.001.

**Figure S2.**
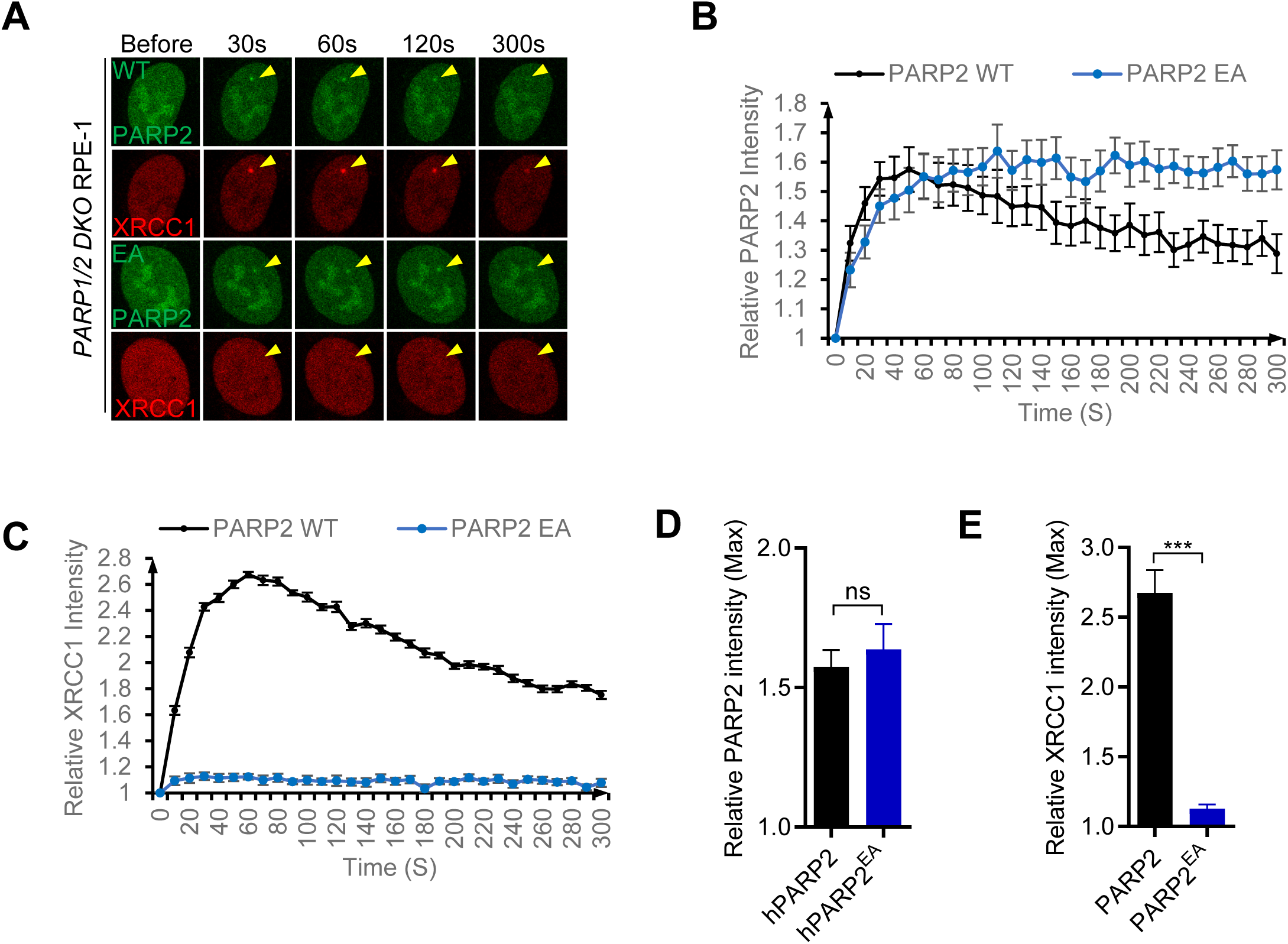
The PARP2 E545A mutation eliminates the formation of PAR chains on PARP2. (A) Representative live-cell images of laser-induced foci of GFP-PARP2 and RFP-XRCC1, or GFP-PARP2^E545A^ and RFP-XRCC1 in *PARP1/2 DKO* RPE-1 cells. (B) The relative intensity kinetics of GFP-PARP2 or GFP-PARP2^E545A^ at DNA damage sites. (C) The relative intensity kinetics for RFP-XRCC1 with GFP-PARP2 or GFP-PARP2^E545A^ at DNA damage sites. (D) The maximal relative intensity of GFP-PARP2 and GFP-PARP2^E545A^ and (E) RFP-XRCC1. The bars and error bars represent means and SEM from one out of three representative experiments with n > 8 cells per experiment. The two-tailed unpaired student’s t-test was used to calculate the p-values. ***p<0.001.

**Figure S3.**
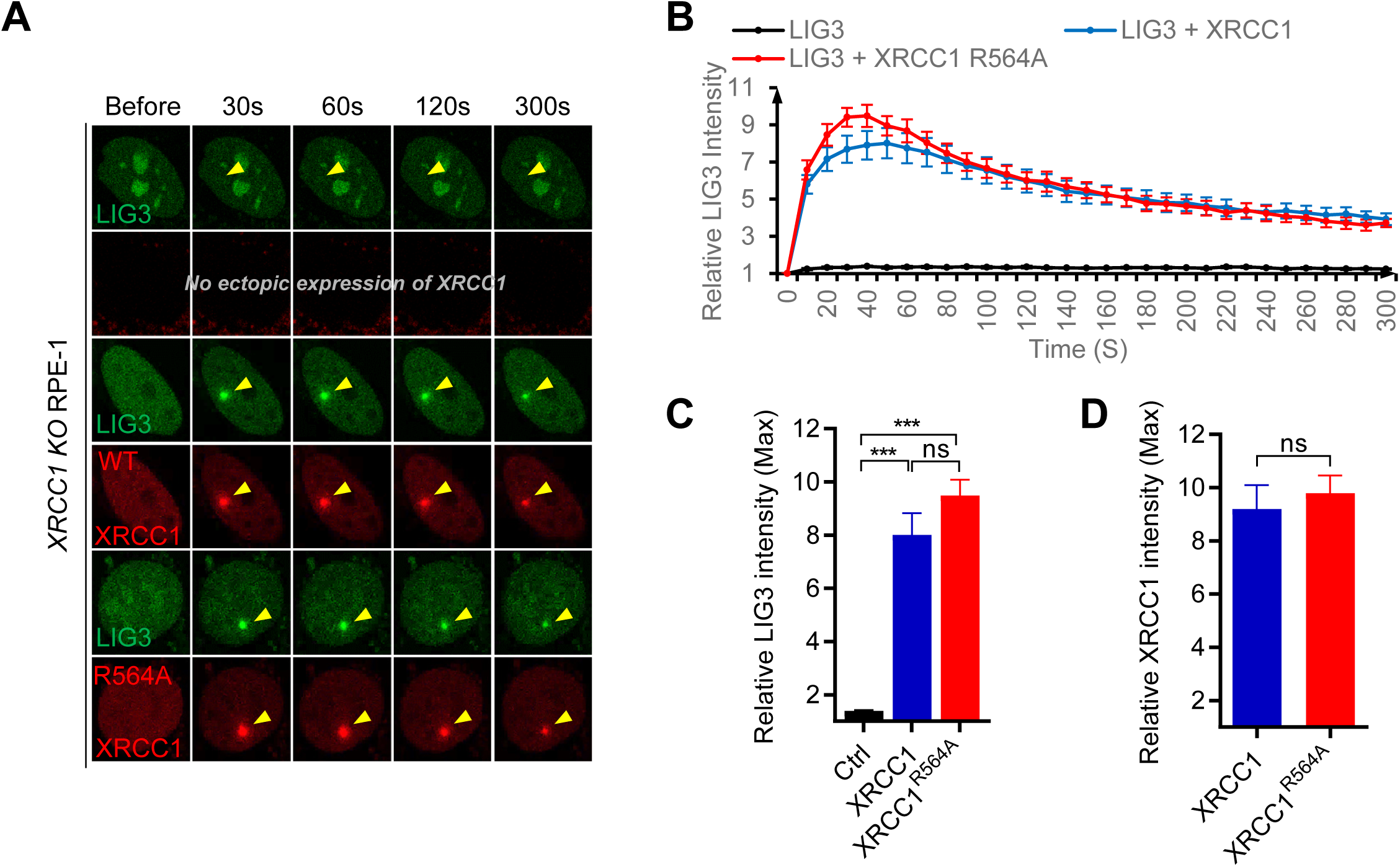
Ectopically expressed XRCC1 R564A can still support Lig3 recruitment. (A) Representative live-cell images of laser-induced foci of GFP-LIG3 alone, GFP-LIG3 and RFP-XRCC1, GFP-LIG3 and RFP-XRCC1^R564A^ in *XRCC1 KO* RPE-1 cells. The yellow arrowheads indicate the areas of micro-irradiation. (B) The relative intensity kinetics for GFP-LIG3 alone or with RFP-XRCC1, or RFP-XRCC1^R564A^ are shown. (C) The maximal relative intensity of GFP-LIG3 and (D) RFP-XRCC1 and RFP-XRCC1^R564A^. The bars and error bars represent means and SEM from one out of two representative experiments with n > 8 cells per experiment. The two-tailed unpaired student’s t-test was used to calculate the p-values. ns, p>0.05, ***p<0.001.

## Notes

### Competing Interest Statement

The authors have declared no competing interest.

